# Effects of B lymphocytes Invitro Treatment with Rose Extract on Transmembrane Ligands

**DOI:** 10.1101/2023.04.13.536644

**Authors:** Mark Christopher Arokiaraj, Eric Menesson

**Affiliations:** Pondicherry Institute of Medical Sciences, Cardiology, Pondicherry, India; Tebu Bio, Lab services, France

## Abstract

**Aims:** The study was performed to evaluate the role of red rose extract (Pierre de Ronsard) on B lymphocytes. Specifically the gene expression of CD20, CD30, CD40, and CCR5 in human B cells were studied after treatment with rose extract.

**Methods:** Red rose extract was prepared at the dilution of 0.0075% (v/v) and stored until use at -20°C. Cell treatment was performed at 37°C on B cells. The cells were plated in 6 well plates at 1.5×10^6^ cells per well and stored at -80°C. Total RNA extraction and quality control were performed. RTq-PCR was performed according to Genecopoeia’s instructions. The cycle threshold method (ΔΔCt) was used for data analysis.

**Results:** The comparative Ct method quantification (2^-ΔCt) and fold change for CD20, CD 30, CD 40 and CCR5 were - 5.65E+01, 4.80E-01, N/A, 2.47E-01; and 0.954,0.377, N/A and 0.577, respectively. The amount of total RNA extracted from about 4.5×10^6^ cells was low and did not allow us to measure the RNA profile. The A260/A230 ratios were very low due to the low amount of RNA. The analysis of gene expression by qRT-PCR showed that CD40 was not expressed in untreated cells and cells treated with rose extract with the Ct values over 32. All other genes were expressed and well-measured in both B cell samples.

**Conclusion:** The treatment with rose extract at 0.0075% (v/v) did not modify the expression of CD20. However, the expression of CD30 and CCR5 decreased with the rose extract treatment. The range of the fold change showed that the result of CD30 expression was more accurate than for CCR5.

## Introduction

B lymphocytes are known for their pivotal role in modulating immunity. They play a significant role in both innate and adaptive immunity are modified by these cells.^1-4^ Autoimmune disorders are modulated by B cells.^2^ B cells have a role in antibody synthesis, antigen presentation, cytokine, and chemokine synthesis.^3^ Cross-reactive antibodies synthesised by B cells help to regulate B cell memory and other immune-modulatory functions.^4^ 9G4 cells function as interfaces between innate and adaptive immunity.^1^ B cell depletion therapies have been used in the treatment for autoimmune disorders.^5^ CD20, CD30, CD 40, and CCR5 ligands are expressed on the surface of the B cells. These ligands are plasma membrane phosphoproteins and have deeper connections in the cells and their expression is modified in various diseases and conditions based on the cell and extracellular signals. CD 20 is expressed in a wide range of tumours and autoimmune disorder regulation.^6-9^

CD30 expression is found on Hodgkin and Reed-Sternberg cells, anaplastic large-cell lymphoma cells, and on activated B or T lymphocytes. CD30 has been shown to be a transmembrane receptor that is significantly homologous to the tumour necrosis factor receptor (TNFR) family.^10,11^ CD40 is expressed primarily by activated T cells, as well as activated B cells and platelets; and under inflammatory conditions is also induced on monocytic cells, natural killer cells, mast cells, and basophils.^12-15^ CCRF motif is multifunctional and plays a role in immune regulation, and neointimal proliferation.^16,17^

Currently, there are many drugs which modulate the effect of these ligands in various diseases.^18-25^ However, these medications are also associated with various side effects profile, which would limit their use.^18-25^ In this study, the rose extract was used for the evaluation on the B cells. The regulation of these ligands on the B cell surface will give more insights in the immune modulation of the rose extract. In our previous study, we have shown that the rose extract at concentrations of 0.5% or less is not associated with cell lysis.^26^

## Methods

Six vials of B cells derived from human PBMC using positive immunomagnetic selection directly against CD19 and cryopreserved immediately after isolation were thawed, pooled, and plated in 6 wells of a 6-well plate at 1.5×10^6^ cells per well. After 24h, the cells of 3 wells were treated with 0.0075% (v/v) rose extract prepared during the project POMC-032018 for 24h. In the other 3 wells, the B cells were untreated.

After 24h incubation, the cells were collected as treated and untreated cells, and washed with PBS twice. They were finally centrifuged and stored at -80°C as cell pellets until RNA extraction.

Total RNA was extracted using NucleoSpin RNA plus kit (Macherey-Nagel, ref. 740984.10) according to the manufacturer’s protocol, including the DNase treatment. Extracted RNA was stored at -80°C for qRT-PCR assay.

The qRT-PCR was conducted according to the Genecopoeia’s instructions. The BlazeTaqTM One-Step SYBR® Green RT-qPCR kit is designed to perform RT and real-time PCR in one step. The qRT-PCR was performed using 6ng of total RNA. Two controls were done per sample, without RTase and without RNA template (NTC). The required volumes for qRT-PCR are presented in table 1.

**Table 1:** Description of qRT-PCR reaction mix.

| Component | Volume for test | Volume for no RTase control | Volume for no RNA template control |
| --- | --- | --- | --- |
| BlazeTaq One Step RT-qPCR Mix (5x) | 4 | 4 | 4 |
| BlazeTaq RTase Mix (50x) | 0.4 | 0 | 0.4 |
| PCR primer mix (2 μM) | 2 | 2 | 2 |
| RNA template (6 ng/reaction) | 5 | 5 | 0 |
| dd H<sub>2</sub>O | 8.6 | 9 | 13.6 |
| **Total Volume** | **20** | **20** | **20** |

The components mix were distributed in PCR plate. Primer mix and RNA template were added to corresponding wells. PCR was performed using Eppendorf Mastercycler RealPlex 2S by following the PCR program presented in table 2.

**Table 2:** Timing of qRT-PCR program.

| Cycles | Step | Temperature | Time |
| --- | --- | --- | --- |
| 1 | Reverse Transcription | 42°C | 10 min |
| 1 | Initial Denaturation | 95°C | 3 min |
| 40 | Denaturation | 95°C | 10 s |
|  | Extension | 60°C | 30 sec |
|  | Melting curve |  |  |

## Results

Absorbance spectra values was first measured by NanoVue (GE Healthcare). The amount of total RNA extracted from about 4.5×10^6^ cells was low and did not allow us to measure the RNA profile by using the TapeStation instrument (Table 3). The A260/A230 ratios were very low, indicating a relatively high proportion of salts in the samples, but the reason was the very low amount of RNA rather than an unexpected quantity of salts after extraction.

**Table 3:**
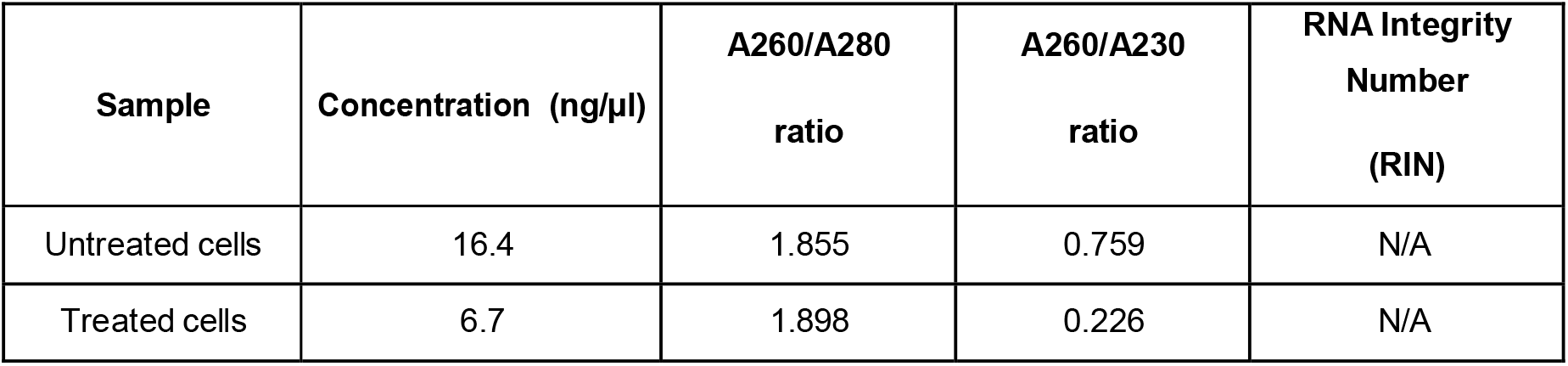
Absorbance of extracted RNAs.

| Sample | Concentration (ng/μl) | A260/A280 ratio | A260/A230 ratio | RNA Integrity Number (RIN) |
| --- | --- | --- | --- | --- |
| Untreated cells | 16.4 | 1.855 | 0.759 | N/A |
| Treated cells | 6.7 | 1.898 | 0.226 | N/A |

The ΔΔCt (cycle threshold) was used for data analysis. In separate reactions, Ct value was determined for each replicate of the housekeeping gene HPRT1 and UBE2D2 (HKG) and gene of interest (GOI) in both samples. For each sample, the difference between the Ct average of GOI replicate and the Ct average for the HKG was calculated (ΔCt). The normalized GOI gene expression was then determined as 2(–ΔΔCt).

The fold-change is the normalized gene expression in the treated sample divided the normalized gene expression in untreated cells. Raw data are listed in table 4 and the normalized gene expressions and fold changes are shown in table 5.

**Table 4:** Ct values from the qRT-PCR.

|  |  | Test |  | no Rtase |  | NTC |
| --- | --- | --- | --- | --- | --- | --- |
|  |  | Untreated | Treated | Untreated | Treated |  |
| HPRT1 | Triplicate 1 | 26.92 | 27.38 | - | - | - |
|  | Triplicate 2 | 27.03 | 27.53 | - | - | - |
|  | Triplicate 3 | 26.67 | 27.50 | - | - | - |
|  | Avg | 26.87 | 27.47 | - | - | - |
| UBE2D2 | Triplicate 1 | 28.68 | 29.20 | - | - | - |
|  | Triplicate 2 | 28.54 | 28.91 | - | 39.79 | - |
|  | Triplicate 3 | 28.67 | 29.03 | - | 36.84 | - |
|  | Avg | 28.63 | 29.04 | - | 38.31 | - |
| CD20 | Triplicate 1 | 21.94 | 22.43 | 37.24 | 38.45 | 33.27 |
|  | Triplicate 2 | 22.06 | 22.79 | 37.72 | 37.98 | 32.86 |
|  | Triplicate 3 | 21.80 | 22.30 | 36.96 | 35.96 | 33.09 |
|  | Avg | 21.93 | 22.51 | 37.30 | 37.46 | 33.07 |
| CD30 | Triplicate 1 | 29.36 | 30.87 | - | - | 34.37 |
|  | Triplicate 2 | 28.35 | 30.69 | - | - | 36.71 |
|  | Triplicate 3 | 28.70 | 30.60 | 38.04 | - | 33.64 |
|  | Avg | 28.81 | 30.72 | - | - | 34.91 |
| CD40 | Triplicate 1 | 34.70 | 40.81 | - | - | 36.48 |
|  | Triplicate 2 | 36.74 | 35.67 | 34.95 | - | 35.27 |
|  | Triplicate 3 | 34.13 | 35.97 | 34.14 | 36.51 | 35.00 |
|  | Avg | 35.19 | 37.48 | 34.55 | - | 35.58 |
| CCR5 | Triplicate 1 | 29.75 | 31.60 | - | - | 34.85 |
|  | Triplicate 2 | 29.69 | 30.82 | - | - | 37.76 |
|  | Triplicate 3 | 29.86 | 30.78 | - | 39.13 | 35.67 |
|  | Avg | 29.77 | 31.07 | - | - | 36.09 |

**Table 5:**
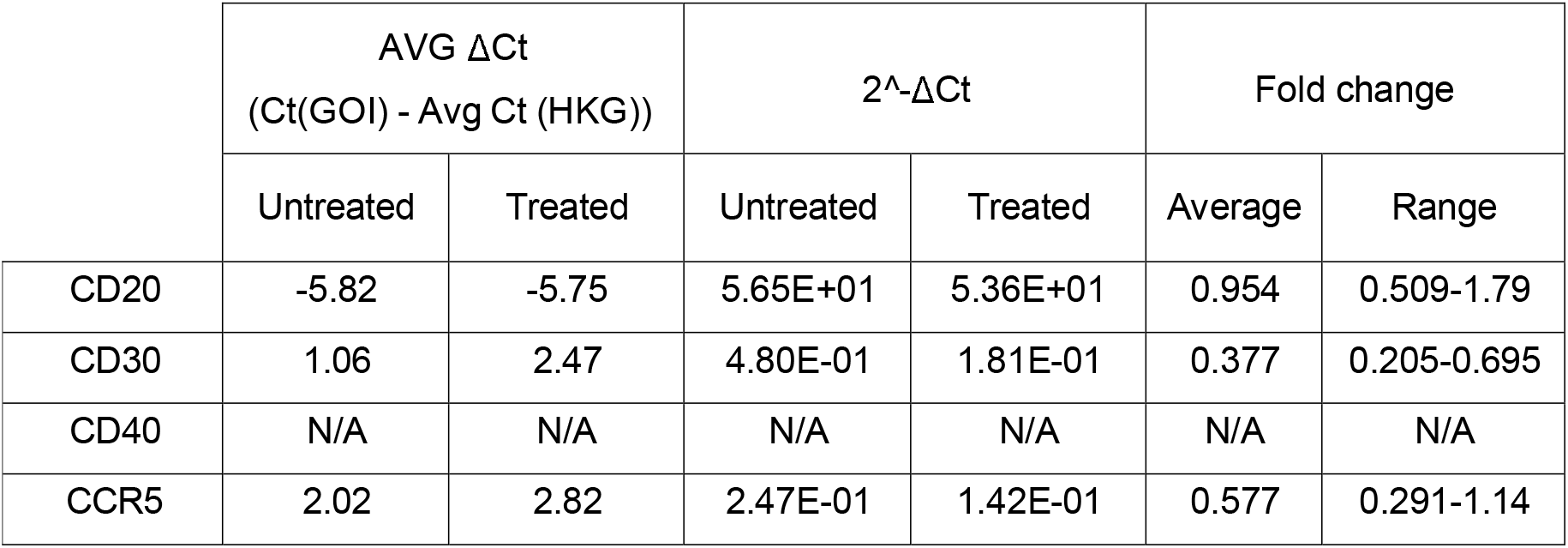
Gene expression fold change between treated and untreated B cells.

In this study, the expression of CD30 on the B cells is reduced after treatment with rose extract (0.0075%). It reduces to 0.377 fold change compared to untreated cells, and the results are consistent (0.205 to 0.695). In our previous studies we have demonstrated that the rose extract at concentrations less than 0.5% is not cytotoxic to the cells. The expression of CCR5 is also reduced by -0.577 fold change, but the reduction is not consistent as it crosses the non-specific mark (0.29 to 1.14).

## Discussion

The analysis of gene expression by qRT-PCR showed that CD40 was not expressed in untreated cells and cells treated with rose extract with the Ct values over 32. The CD20 levels were not significantly altered by the rose extract treatment in this study. All other genes were expressed and were well measured in both B cell treated and untreated cell samples.

The results could translate to clinical benefits by the reduction in CD30 expression in cancerous cells. CD30 is an active marker in major tumours like Hodgkin’s lymphoma, anaplastic large cell lymphoma. ^27, 28^ It also plays a role in angiogenic cancers like angiosarcomas, epithelioid hemangioendothelioma, Kaposi’s sarcoma etc. ^29^

Also, CD30 inhibition is associated with the treatment of primary effusion lymphoma. ^30^ CD30 is also an active marker in autoimmune disorders but its role in this treatment is not known in-depth. Targeting CD 30-30L has been used in the treatment of autoimmune and inflammatory disorders.^31^ Remission of rheumatoid arthritis has been observed in the treatment with anti-CD30 agent, brentuximab vedotin.^32^ Inhibition of CD30 in T cells is also associated with reduced graft vs host disease,^33^ and immune complex-mediated glomerulonephritis.^34^ CD30-30L interaction inhibition is associated with reduced atherosclerosis in a bench study.^35^ Soluble CD30 is reduced in cases with stable coronary artery disease.^36^ CD30 expression is increased in patients with chronic obstructive pulmonary disease and in vascular remodelling.^37^

Similarly, the efficacy of the rose extract in acquired immunodeficiency disorder treatment by the reduction in CCR5 on the T cell surface needs to be studied in retrovirus-infected cells. In our previous study, we have shown the effects in the reduction of soluble CCR5 in endothelial cells treated with rose extract.^17^ CCR5 has also been shown in the regulation of vascular, neurological, and autoimmune disorders signalling.^38^ CCR5 is also involved in learning, memory, and cognitive functions.^39^

Rose extract has been known for its anti-inflammatory, antioxidant, antidiabetic, and antidepressant effects.^40, 41^ It has certain antibacterial activity also.^40, 42^ In our previous studies also, we have shown anti-inflammatory effects of rose extract on T cells. This study shows the beneficial effects by a reduction in CD30 levels, and CCR5 levels on the B cell surface to some extent.

### Limitations

The study was done *invitro*. Large studies are required involving more sample size is required to evaluate the concept in *invitro* and *in vivo*. The study was very focused on the cell certain ligands on the cell surface. Further evaluation is required with the detailed study on cytokines for more information on the B cell function after 0.0075% rose extract treatment.

## Conclusions

The analysis of gene expression by qRT-PCR showed that CD40 was not expressed in untreated cells and cells treated with rose extract with Ct values over 32. All other genes were expressed and well-measured in both B cell samples. The treatment with rose extract at 0.0075% (v/v) did not modify the expression of CD20. However, the expression of CD30 and CCR5 seemed to be decreased by the treatment. The range of the fold change showed that the result of CD30 expression was more accurate than for CCR5.

## Ethical considerations

Not applicable

## Author contributions

MCA conceived the idea and method and wrote the paper, EM performed the experiments and derived the results.

## Bibliography

1. Milner EC, Anolik J, Cappione A, Sanz I. Human innate B cells: a link between host defense and autoimmunity? Springer Semin Immunopathol. 2005 Mar;26(4):433–52. doi: 10.1007/s00281-004-0188-9. Epub 2005 Jan 5. PMID: 15633016; PMCID: PMC1431976.

2. Tsay GJ and Zouali M (2018) The Interplay Between Innate-Like B Cells and Other Cell Types in Autoimmunity. Front. Immunol. 9:1064. doi: 10.3389/fimmu.2018.01064

3. Zhang, X. Regulatory functions of innate-like B cells. Cell Mol Immunol 10, 113–121 (2013). https://doi.org/10.1038/cmi.2012.63

4. Grasseau, A., Boudigou, M., Le Pottier, L. et al. Innate B Cells: the Archetype of Protective Immune Cells. Clinic Rev Allerg Immunol 58, 92–106 (2020). https://doi.org/10.1007/s12016-019-08748-7.

5. Lee, D.S.W., Rojas, O.L. & Gommerman, J.L. B cell depletion therapies in autoimmune disease: advances and mechanistic insights. Nat Rev Drug Discov 20, 179–199 (2021). https://doi.org/10.1038/s41573-020-00092-2

6. Pavlasova G, Mraz M. The regulation and function of CD20: an “enigma” of Bcell biology and targeted therapy. Haematologica. 2020 Jun;105(6):1494–1506. doi: 10.3324/haematol.2019.243543. PMID: 32482755; PMCID: PMC7271567.

7. Kläsener K, Jellusova J, Andrieux G, Salzer U, Böhler C, Steiner SN, et al. CD20 as a gatekeeper of the resting state of human B cells. Proceedings of the National Academy of Sciences. 2021;118(7). 10.1073/pnas.202134211

8. https://www.pathologyoutlines.com/topic/cdmarkerscd20.html

9. Michot JM, Buet-Elfassy A, Annereau M, Lazarovici J, Danu A, Sarkozy C, Chahine C, Bigenwald C, Bosq J, Rossignol J, Romano-Martin P, Baldini C, Ghez D, Dartigues P, Massard C, Ribrag V. Clinical significance of the loss of CD20 antigen on tumor cells in patients with relapsed or refractory follicular lymphoma. Cancer Drug Resistance. 2021; 4(3):710–8. http://dx.doi.org/10.20517/cdr.2020.109

10. de Bruin PC, Gruss HJ, van der Valk P, Willemze R, Meijer CJ. CD30 expression in normal and neoplastic lymphoid tissue: biological aspects and clinical implications. Leukemia. 1995 Oct;9(10):1620–7. PMID: 7564499.

11. van der Weyden, C., Pileri, S., Feldman, A. et al. Understanding CD30 biology and therapeutic targeting: a historical perspective providing insight into future directions.Blood Cancer J. 7, e603 (2017). https://doi.org/10.1038/bcj.2017.85

12. Funakoshi S, Longo DL, Beckwith M, Conley DK, Tsarfaty G, Tsarfaty I, et al. Inhibition of human B-cell lymphoma growth by CD40 stimulation. Blood. 1994;83(10):2787–94.

13. Linderoth J, Ehinger M, Jerkeman M, Bendahl P-O, Åkerman M, Berglund M, et al. CD40 expression identifies a prognostically favourable subgroup of diffuse large B-cell lymphoma. Leukemia & Lymphoma. 2007;48(9):1774–9.

14. Feng Z, Wang J. Soluble CD40 ligand inhibits the growth of non-hodgkin’s lymphoma cells through the JNK signaling pathway. Oncology Letters. 2020;21(1).

15. Karnell JL, Rieder SA, Ettinger R, Kolbeck R. Targeting the CD40-CD40L pathway in autoimmune diseases: Humoral immunity and beyond. Adv Drug Deliv Rev. 2019 Feb 15;141:92–103. doi:10.1016/j.addr.2018.12.005. Epub 2018 Dec 13. PMID: 30552917.

16. Oppermann M. Chemokine receptor CCR5: insights into structure, function, and regulation. Cell Signal. 2004 Nov;16(11):1201–10. doi: 10.1016/j.cellsig.2004.04.007. PMID: 15337520.

17. Arokiaraj, M., & Menesson, E. (2021). Effect of rose extract treatment on soluble CCR5 and CXCR4 secretion by the endothelial cells invitro. Biomedical Research and Therapy, 8(5), 4333–4344. https://doi.org/10.15419/bmrat.v8i5.672

18. Hansel, T., Kropshofer, H., Singer, T. et al. The safety and side effects of monoclonal antibodies. Nat Rev Drug Discov 9, 325–338 (2010).

19. Du FH, Mills EA, Mao-Draayer Y. Next-generation anti-CD20 monoclonal antibodies in autoimmune disease treatment. Auto Immun Highlights. 2017 Nov 16;8(1):12. doi: 10.1007/s13317-017-0100-y. PMID: 29143151; PMCID: PMC5688039.

20. Payandeh Z, Bahrami AA, Hoseinpoor R, Mortazavi Y, Rajabibazl M, Rahimpour A, et al. The applications of anti-cd20 antibodies to treat various B cells disorders. Biomedicine & Pharmacotherapy. 2019;109:2415–26.https://doi.org/10.1016/j.biopha.2018.11.121

21. van Vollenhoven RF, Emery P, Bingham CO, Keystone EC, Fleischmann RM, Furst DE, et al. Long-term safety of rituximab in rheumatoid arthritis: 9.5-year follow-up of the global clinical trial programme with a focus on adverse events of interest in RA patients. Annals of the Rheumatic Diseases. 2012;72(9):1496–502. http://dx.doi.org/10.1136/annrheumdis-2012-201956.

22. Yi JH, Kim SJ, Kim WS. Brentuximab Vedotin: Clinical updates and practical guidance. Blood Research. 2017;52(4):243.

23. Wang, Y., Nowakowski, G.S., Wang, M.L. et al. Advances in CD30- and PD-1-targeted therapies for classical Hodgkin lymphoma. J Hematol Oncol 11, 57 (2018). https://doi.org/10.1186/s13045-018-0601-9

24. Ramos CA, Grover NS, Beaven AW, Lulla PD, Wu M-F, Ivanova A, et al. Anticd30 car-T cell therapy in relapsed and refractory Hodgkin lymphoma. Journal of Clinical Oncology. 2020;38(32):3794–804.

25. Muta H, Podack ER. CD30: from basic research to cancer therapy. Immunol Res. 2013 Dec;57(1-3):151–8. doi: 10.1007/s12026-013-8464-1. PMID: 24233555; PMCID: PMC3992519.

26. Arokiaraj MC, Menesson E.Novel anti-inflammatory and immunomodulation effects of rose on the endothelium in normal and hypoxic invitro conditions. Angiol Cir Vasc. 2020 Feb. 5;15(4):238–4.

27. Muta H, Podack ER. CD30: from basic research to cancer therapy. Immunol Res. 2013 Dec;57(1-3):151–8. doi: 10.1007/s12026-013-8464-1. PMID: 24233555; PMCID: PMC3992519.

28. CD30 [Internet]. Pathology Outlines - PathologyOutlines.com. Available from: https://www.pathologyoutlines.com/topic/cdmarkerscd30.html

29. Alimchandani M, Wang ZF, Miettinen M. CD30 expression in malignant vascular tumors and its diagnostic and clinical implications: a study of 146 cases. Appl Immunohistochem Mol Morphol. 2014 May-Jun;22(5):358–62. doi: 10.1097/PAI.0000000000000048. PMID: 24805132; PMCID: PMC4017247.

30. CD30 targeting with brentuximab vedotin: a novel therapeutic approach to primary effusion lymphoma. https://doi.org/10.1182/blood-2013-01-481713

31. Oflazoglu E, Grewal IS, Gerber H. Targeting CD30/CD30L in oncology and autoimmune and inflammatory diseases. Adv Exp Med Biol. 2009;647:174–85. doi: 10.1007/978-0-387-89520-8_12. PMID: 19760074.

32. Pankit Vachhani, Nilanjana Bose, James P. Brodeur, Beata Holkova, Prithviraj Bose, Remission of rheumatoid arthritis on brentuximab vedotin, Rheumatology, Volume 53, Issue 12, December 2014, Pages 2314–2315, https://doi.org/10.1093/rheumatology/keu374

33. Chen YB, McDonough S, Hasserjian R, Chen H, Coughlin E, Illiano C, Park IS, Jagasia M, Spitzer TR, Cutler CS, Soiffer RJ, Ritz J. Expression of CD30 in patients with acute graft-versus-host disease. Blood. 2012 Jul 19;120(3):691–6. doi: 10.1182/blood-2012-03-415422. Epub 2012 Jun 1. PMID: 22661699; PMCID: PMC3401221.

34. Artinger K, Kirsch AH, Mooslechner AA, Cooper DJ, Aringer I, Schuller M, et al. Blockade of tumor necrosis factor superfamily members CD30 and OX40 abrogates disease activity in murine immune-mediated glomerulonephritis. Kidney International. 2021;100(2):33648. https://doi.org/10.1016/j.kint.2021.02.039

35. Interference of the CD30–CD30L Pathway Reduces Atherosclerosis Development. Arteriosclerosis, Thrombosis, and Vascular Biology. 2012;32:2862–2868. https://doi.org/10.1161/ATVBAHA.112.300509.

36. Mahmoudi MJ, Hedayat M, Rezaei M, Yaraghi AS, Mahmoudi M. In vitro Soluble CD30 Levels in Patients with Chronic Stable Coronary Artery Disease. Iran Jrnl Allergy asthma Immunol.2011; 10(4): 237 242.

37. Luo L, Liu Y, Chen D, Chen F, Lan HB, Xie C. CD30 is highly expressed in chronic obstructive pulmonary disease and induces the pulmonary vascular remodeling. BioMed Research International. 2018;2018:1–13.

38. Jones KL, Maguire JJ, Davenport AP. Chemokine receptor CCR5: from AIDS to atherosclerosis. Br J Pharmacol. 2011 Apr;162(7):1453–69. doi: 10.1111/j.1476-5381.2010.01147.x. PMID: 21133894; PMCID: PMC3057285.

39. Necula D, Riviere-Cazaux C, Shen Y, Zhou M. Insight into the roles of CCR5 in learning and memory in normal and disordered states. Brain, Behavior, and Immunity. 2021;92:1–9.

40. Mahboubi M. Rosa Damascena as holy ancient herb with novel applications. Journal of Traditional and Complementary Medicine. 2016;6(1):10–6.

41. Lee M-hee, Nam TG, Lee I, Shin EJ, Han A-ram, Lee P, et al. Skin antiinflammatory activity of Rose petal extract (rosa gallica) through reduction of MAPK signaling pathway. Food Science & Nutrition. 2018;6(8):2560–7.

42. Yusra Safdar and Taqdees Malik. Antibacterial activity of rose extract. Open Acess journal of complementary and alternative medicine, 2020; 2(4).

